# Liquid-liquid phase separation and fibrillization of tau are independent processes with overlapping conditions

**DOI:** 10.1101/702126

**Authors:** Yanxian Lin, Yann Fichou, Zhikai Zeng, Nicole Y. Hu, Songi Han

## Abstract

Amyloid aggregation of the microtubule binding protein tau is a hallmark of Alzheimer’s disease and many other neurodegenerative diseases. Recently, tau has been found to undergo liquid-liquid phase separation (LLPS) near physiological conditions. Although LLPS and aggregation have been shown to simultaneously occur under certain common conditions, it remains to be seen whether tau LLPS promotes aggregation, or if they are two independent processes. In this study, we address this question by combining multiple biochemical and biophysical assays *in vitro*. We investigated the impacts of LLPS on tau aggregation at three stages: conformation of tau, kinetics of aggregation and fibril quantity. We showed that none of these properties are influenced directly by LLPS, while amyloid aggregation propensity of tau can be altered without affecting its LLPS behavior. LLPS and amyloid aggregation of tau occur under overlapping conditions of enhanced intermolecular interactions and localization, but are two independent processes.

## Introduction

Tau is a protein abundant in neurons of the central nervous system that binds and stabilizes microtubules [1]–[3]. In healthy conditions, tau is highly hydrophilic and soluble in aqueous environment under wide ranges of pH, temperature and ionic strength. However, under pathological conditions, tau forms irreversible and insoluble amyloid aggregates. Amyloid aggregation of tau into specific fibril strain is a hallmark of Alzheimer’s disease (AD) [4], [5], and has been observed under several other neurodegenerative diseases including progressive supranuclear palsy, frontotemporal dementia, Pick’s disease and chronic traumatic encephalopathy [6], [7]. Despite the pathological significance of tau amyloid aggregation, the pathways from soluble tau to insoluble amyloid aggregates have still not been identified nor replicated in any non-human system, neither *in vitro* nor *in vivo*. Finding the pathologically relevant amyloid aggregation pathway, as manifested in the generation of disease phenotypic fibril structures [8]–[10] is critical for understanding the underlying disease pathogenesis and for developing therapies.

Liquid-liquid phase separation (LLPS) is a process that results from multi-valent weak interactions, preferably between flexible polymers [11]–[14]. LLPS yields a polymer-dilute and polymer-rich phase, where the latter is often observed in the form of droplets and retains liquid-like mobility [15]–[20]. Such process has been found to be essential for multiple cellular functions for the last 10 years [13], [21], [22]. However, its association with amyloid aggregation has only recently become a focus [23]–[27], with the LLPS of tau discovered and discussed only recently [28]–[34].

A recent study from our group and collaborators found that tau binds RNA *in vivo* and undergoes liquid-liquid phase separation (LLPS) *in vitro* under a range of polymer concentration, temperature, ionic strength and stoichiometry between RNA and tau [30]. We further mapped out its phase diagram by experiment and field-theoretic simulation. The phase diagram revealed a narrow equilibrium window near cellular conditions, demonstrating that cells can readily access conditions to assemble and dissolve tau-RNA LLPS *in vivo* [31]. Meanwhile, Ambadipudi *et al* in 2017 [28] and Wegmann *et al* in 2018 [29] reported that tau can also undergo LLPS without RNA or other reactants under physiological conditions. In these studies, the conditions for LLPS coincide with those for amyloid aggregation, and therefore LLPS was postulated to mediate and facilitate aggregation. Despite the observation that tau LLPS and amyloid formation occur in sequential order, it is unclear whether LLPS facilitates aggregation of tau or if both processes occur independently, but merely have overlapping conditions. This study sets out to unravel whether LLPS influences the aggregation of tau and *vice versa*.

## Results

### Tau amyloid aggregation colocalizes with LLPS

We used truncated variants of the longest human tau isoform, 2N4R, that contains four repeat domains (R1 to R4) and the entire C-terminal region (residues 255-441, named tau187 here [35]) to study tau-polyanion liquid-liquid phase separation (LLPS) and amyloid aggregation. Four variants of tau187 were used in this work, as listed in (Table 1) and summarized here: 1) tauS: tau187 with C291S mutation; 2) tauSS: tau187 with C291S and C322S mutations; 3) tauSP301L: tauS with additional P301L mutation; 4) tauSSP301L: tauSS with additional P301L mutation. At neutral pH, all four variants are positively charged with an estimated +11 net charge per tau molecule, while full length 2N4R tau is estimated to have a +3 net positive charge per tau molecule (Table 1).

**Table 1.**
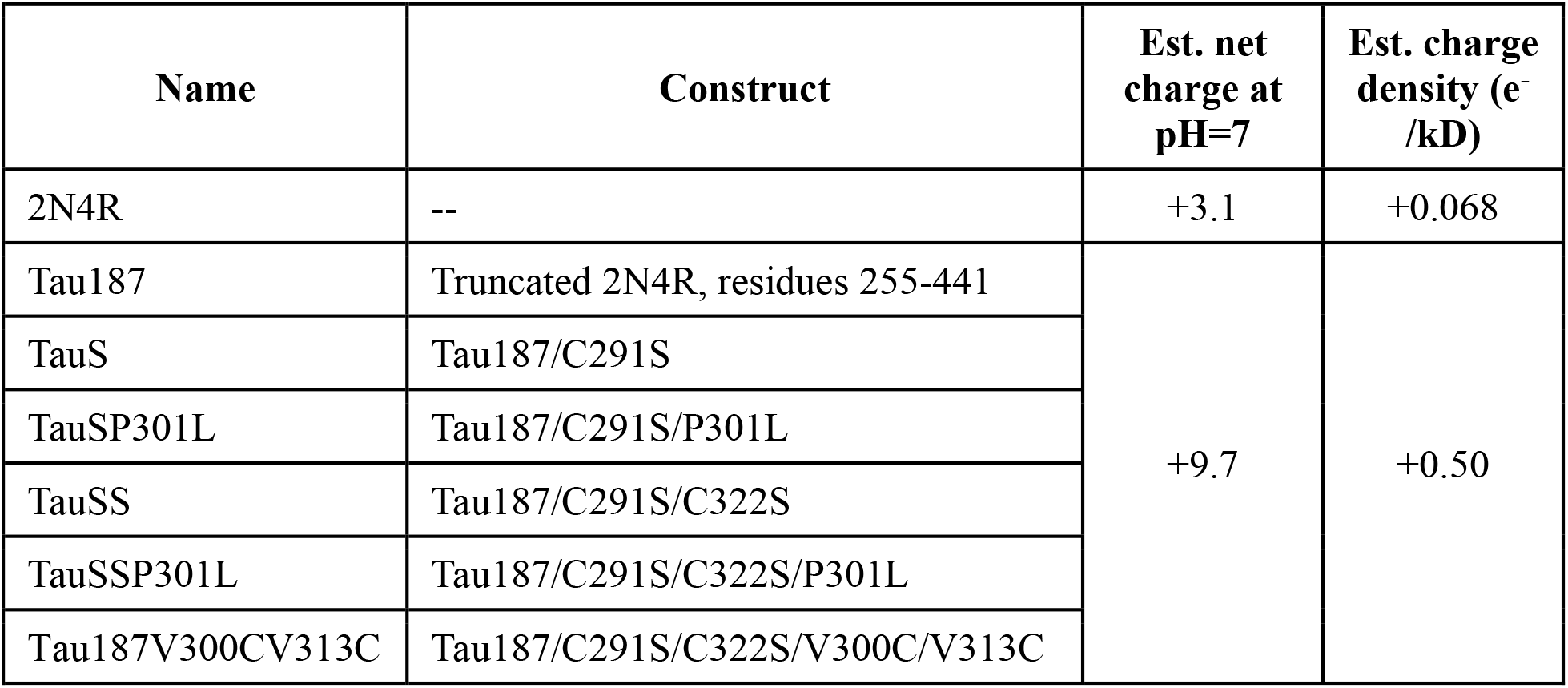
Names and constructs of different tau variants. Full length longest isoform of human tau, 2NR4, is listed with truncated version, tau187, as well as its mutated variants. The net charge at pH = 7 is estimated by Innovagen’s Peptide Property Calculator (http://pepcalc.com/), using primary sequence as input. The estimated charge density is calculated using net charge at pH = 7 divided by molecular weight in kD.

| Name | Construct | Est. net charge at pH=7 | Est. charge density (e<sup>-</sup>/kD) |
| --- | --- | --- | --- |
| 2N4R | -- | +3.1 | +0.068 |
| Tau187 | Truncated 2N4R, residues 255-441 | +9.7 | +0.50 |
| TauS | Tau187/C291S |  |  |
| TauSP301L | Tau187/C291S/P301L |  |  |
| TauSS | Tau187/C291S/C322S |  |  |
| TauSSP301L | Tau187/C291S/C322S/P301L |  |  |
| Tau187V300CV313C | Tau187/C291S/C322S/V300C/V313C |  |  |

Tau has been reported to undergo LLPS with RNA under certain conditions. However, whether this process is driven by tau-RNA specific association or is general to tau-polyanion has not been explicitly tested. To answer this question, tauS was mixed with various polyanions: poly(A) RNA, poly(dA) DNA, heparin and hyaluronic acid (Table 2). The concentrations were chosen so that polyanion and tau reach approximately charge neutrality, i.e., a charge ratio, R, of polyanion:tau of ~1. Immediately after mixing, the clear solution became turbid within seconds, and when imaged under bright field microscope, liquid condensates were visible (Figure 1A). They are characterized by clear round boundaries and merging with each other in seconds. Hence, we refer to these as droplets. Despite having completely different backbone and charged group chemistry, these polyanions can all form droplets upon mixing with tau, suggesting LLPS can be relatively independent of chemical identity, but rather rely on charge matching.

**Table 2.**
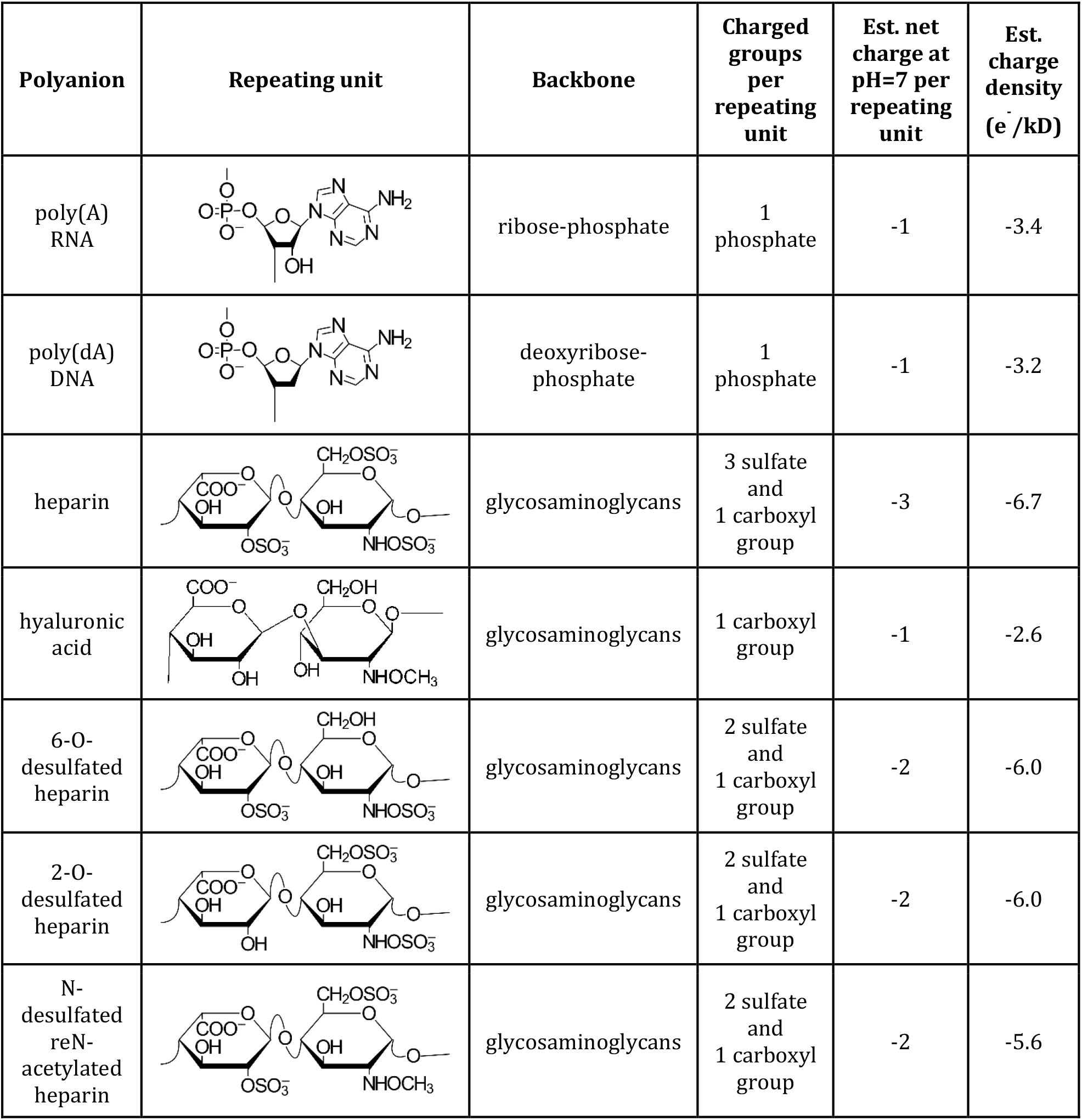
Backbone and charged groups of polyanions. The repeating units for heparin and desulfated heparins represent >70% of the chemical. Estimated charge densities are based on the formula weight of the repeating unit.

| Polyanion | Repeating unit | Backbone | Charged groups per repeating unit | Est. net charge at pH=7 per repeating unit | Est. charge density (e<sup>-</sup>/kD) |
| --- | --- | --- | --- | --- | --- |
| poly(A) RNA |  | ribose-phosphate | 1 phosphate | -1 | -3.4 |
| poly(dA) DNA |  | deoxyribose-phosphate | 1 phosphate | -1 | -3.2 |
| heparin |  | glycosaminoglycans | 3 sulfate and 1 carboxyl group | -3 | -6.7 |
| hyaluronic acid |  | glycosaminoglycans | 1 carboxyl group | -1 | -2.6 |
| 6-O-desulfated heparin |  | glycosaminoglycans | 2 sulfate and 1 carboxyl group | -2 | -6.0 |
| 2-O-desulfated heparin |  | glycosaminoglycans | 2 sulfate and 1 carboxyl group | -2 | -6.0 |
| N-desulfated reN-acetylated heparin |  | glycosaminoglycans | 2 sulfate and 1 carboxyl group | -2 | -5.6 |

**Figure 1.**
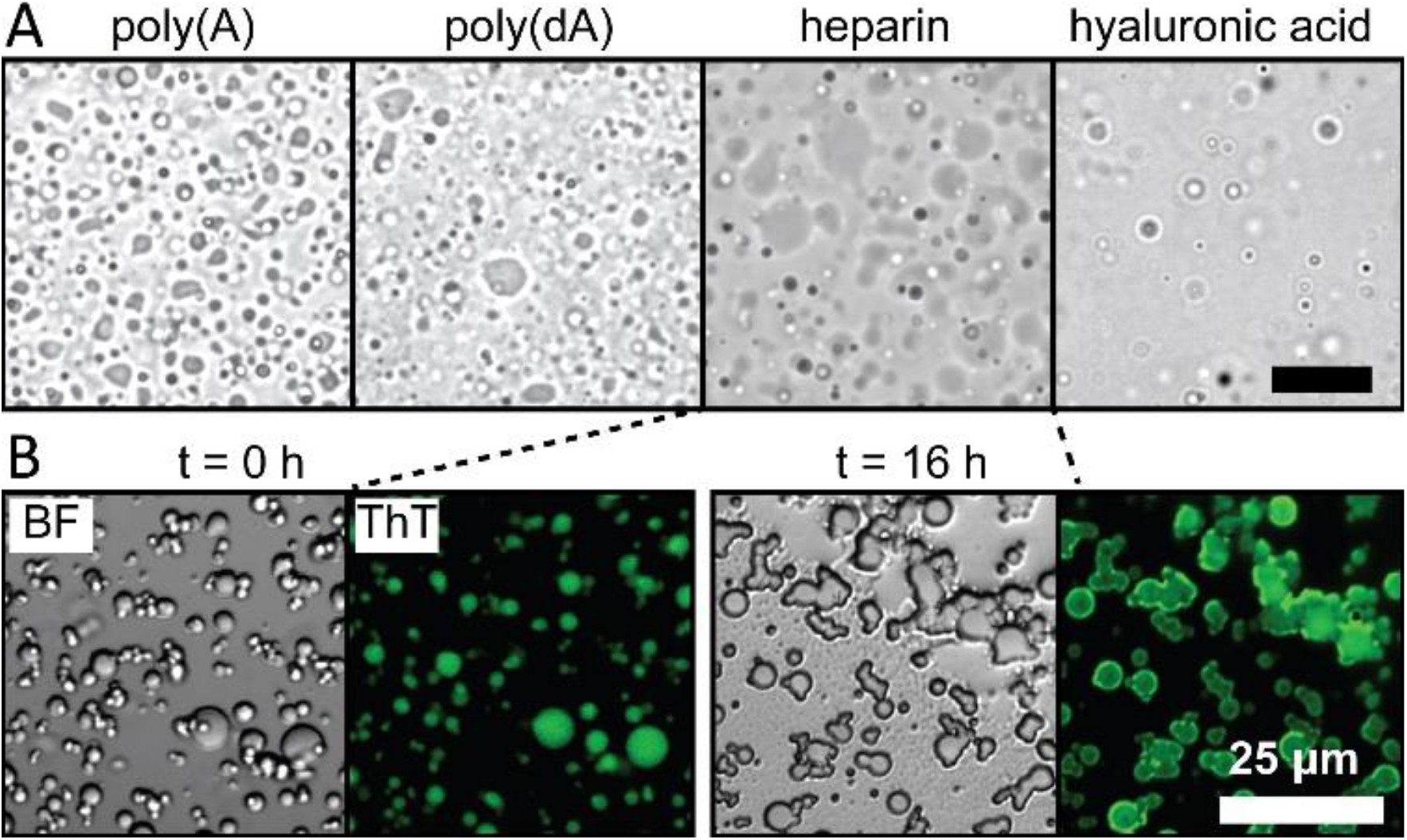
Bright field and confocal microscope images of tau-polyanion LLPS and amyloid aggregation. **A.** 100 μM tauS was mixed with 300 μg/mL poly(A) RNA, 300 μg/mL poly(dA) DNA, 170 μg/mL heparin, 440 μg/mL hyaluronic acid, respectively. Images were taken at room temperature. **B.** Confocal images of tau-heparin LLPS. 100 μM tauS was incubated with 170 μg/mL heparin and 10 μM ThT at 37 °C for 16 h. Confocal images of both bright field (BF) and fluorescence of ThT (λ_emission_=485 nm) were taken immediately after mixing (t = 0h), and 16 hours after incubation (t = 16 h). Scale bars in both A and B are 25 μm long.

We incubated tauSS with heparin at a mixture that achieves net charge neutrality. Confocal microscopy showed that droplets emerged and are retained after 16 hours immediately after mixing (Figure 1B), as well as the localization of amyloid aggregates within the droplet phase. All droplets showed strong fluorescence of ThT, with the fluorescence intensity increase after overnight incubation. Replicas performed at room temperature consistently showed colocalization of ThT fluorescence within droplets (Figure S1). These results demonstrated that tau can undergo amyloid aggregation under conditions identical to that for LLPS, and that the resulting amyloid aggregates colocalize within the dense droplets.

### The extent of amyloid aggregation is independent of LLPS

Despite the observation that amyloid aggregates colocalized within tau droplets, it is not clear whether this is simply because tau is present inside the droplets, or LLPS facilitates aggregation, for instance, through favorable local physical/chemical properties. In the following three sections, we explore whether or not the LLPS state changes tau conformation before aggregation, changes kinetics during aggregation and alters the extent of amyloid fibril formation. We present these questions in reverse order, by first asking whether LLPS influences the quantity of amyloid fibrils formed.

We first investigated the phase behavior of tau-heparin LLPS, by titrating heparin against tauS, and monitoring turbidity reading. The results showed that the turbidity peaks at a charge ratio *R* = 1 for heparin:tau (Figure S2A). At such a condition we prepared another sample and varied the NaCl concentration and monitored turbidity reading. Results showed that the turbidity steadily decreased with increasing [NaCl] to a baseline at [NaCl] > 40 mM (Figure S2B). The systematic effect of ionic strength in modulating intermolecular association is a signature of polyelectrolyte complex coacervation—the mechanism identified for the LLPS of tau and RNA [31].

We next compared LLPS and amyloid aggregation of tau-heparin at varying stoichiometry and ionic strength. We incubated 20 μM of tauSS with heparin at room temperature at neutral pH overnight on a microplate. We varied [NaCl] from 0 to 320 mM on each column and varied [heparin] from *R* = 0 to *R* = 16 on each row. Turbidity and ThT fluorescence of each sample were monitored (Figure S3). The average turbidity reading during the first 1 hour incubation was used to represent the relative amount of droplet, while the maximum ThT fluorescence, extracted from fitting a sigmoidal curve to data (see Materials and Methods), was used to represent the relative amount of amyloid aggregates. Both turbidity and maximum ThT fluorescence were scaled 0 to 1 and presented in a scatter plot side by side (Figure 2). A bivariate normal distribution was fitted and shown as red contour (see Materials and Methods), indicating the condition preferring amyloid aggregation and LLPS. The results showed that amyloid aggregation occurs at a much broader range of [NaCl] and [heparin], while LLPS occurs at a small subset of these conditions (Figure 2 and Figure S3). The fitting results showed that the maximum position for amyloid aggregation is distinct from that for LLPS, showing optimal LLPS is neither necessary nor sufficient for optimal amyloid aggregation. To answer whether this observation is limited to cysteine-free tau, we repeated it using tauS-heparin under non-reducing condition to encourage disulfide bonding. The results showed again that amyloid aggregation occurs under a broader range of [NaCl] and [heparin] at different optimal stoichiometry (Figure S4). We can conclude that the extent of amyloid aggregation is independent of LLPS.

**Figure 2.**
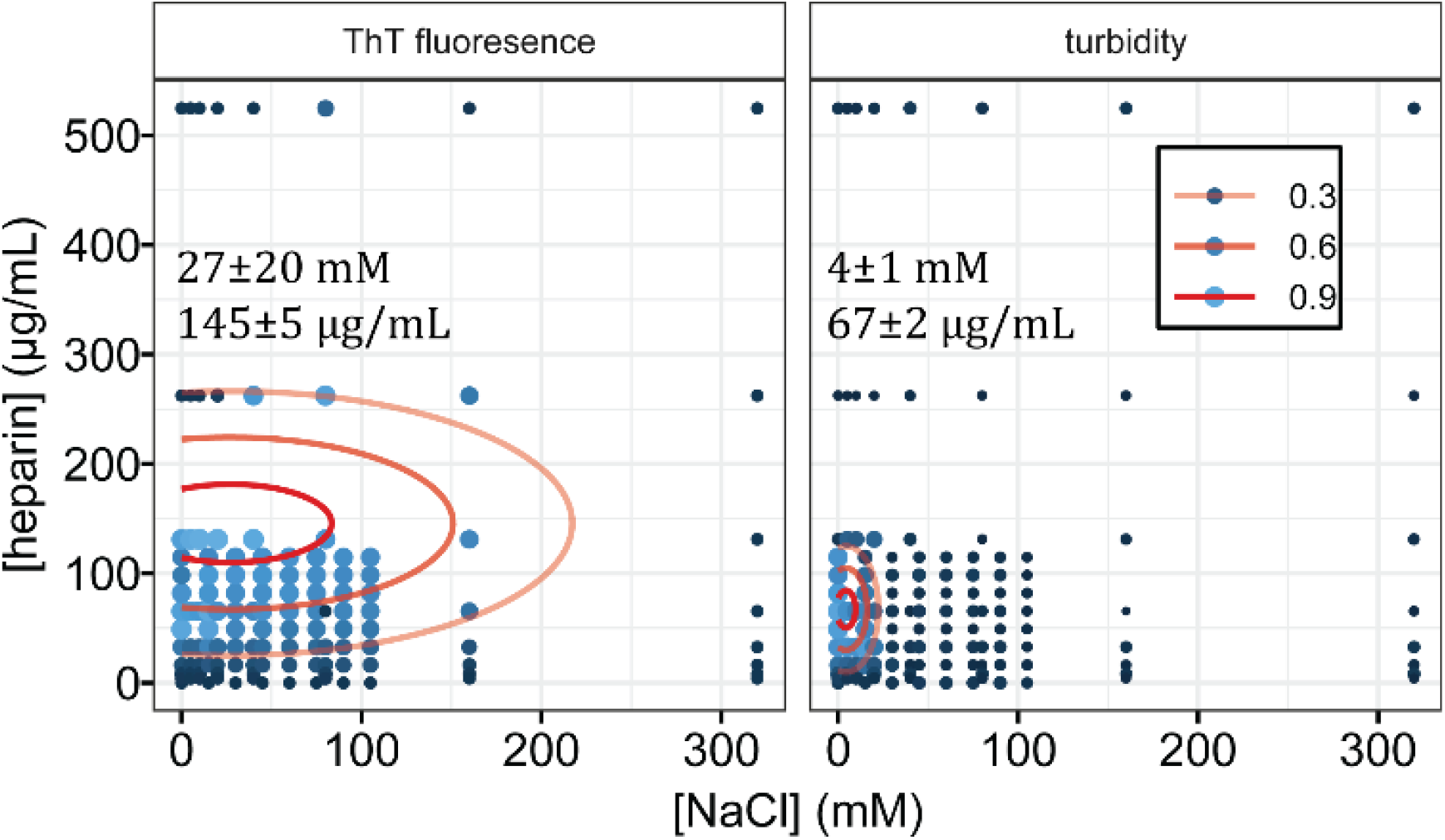
LLPS and amyloid aggregation of tau-heparin at varying [NaCl] and [heparin]. 20 μM of tauSS was incubated with heparin at varying [heparin] from 0 to 544 μg/mL and [NaCl] from 0 to 320 mM. Turbidity and ThT fluorescence of each sample were monitored and the maximum readings were extracted. Both the maximum turbidity and maximum ThT fluorescence across all the samples were normalized to scale of 0~1. The size and color of data points showed the normalized values. Three red solid contour lines from outward to inward showed the fitted bivariate normal distribution at 0.3, 0.6 and 0.9 respectively. Average and standard error of the optimal conditions were listed.

### Half time and monomer dependence of amyloid aggregation are independent of LLPS

LLPS results in the formation of concentrated and viscous droplets of tau and polyanions. As high protein concentration tends to accelerate aggregation, while high viscosity tends to decelerate assembly, it is not immediately obvious whether aggregation of tau in LLPS droplets will slow down or speed up. TauSS-heparin droplets were prepared and mixed with varying [NaCl] and ThT fluorescence and turbidity reading were analyzed. Turbidity of samples were normalized to 1. For qualitative estimation, we regarded conditions where turbidity falls above 0.3 as droplet conditions, and below 0.3 as non-droplet conditions. To quantify the kinetics of tau aggregation, we used the time when ThT fluorescence reading reaches half maximum, referred to as half time or t_1/2_ (see Materials and Methods).

We first investigated t_1/2_ while keeping [heparin] at R = 1 and R = 3 (Figure **3**A). Results showed that overall t_1/2_ increases with increasing [NaCl], regardless of [heparin]. Importantly, near the phase separation boundary, the t_1/2_ value under droplet condition was not significantly different compared to the non-droplet condition (Figure **3**A). Independent repeats of the same conditions in (Figure **3**A) showed similar results (Figure S5B). Meanwhile, the dependence of t_1/2_ on [heparin] from R = 0.5 to R = 4 was also investigated, and shown to be independent of LLPS formation (Figure **3**B).

**Figure 3.**
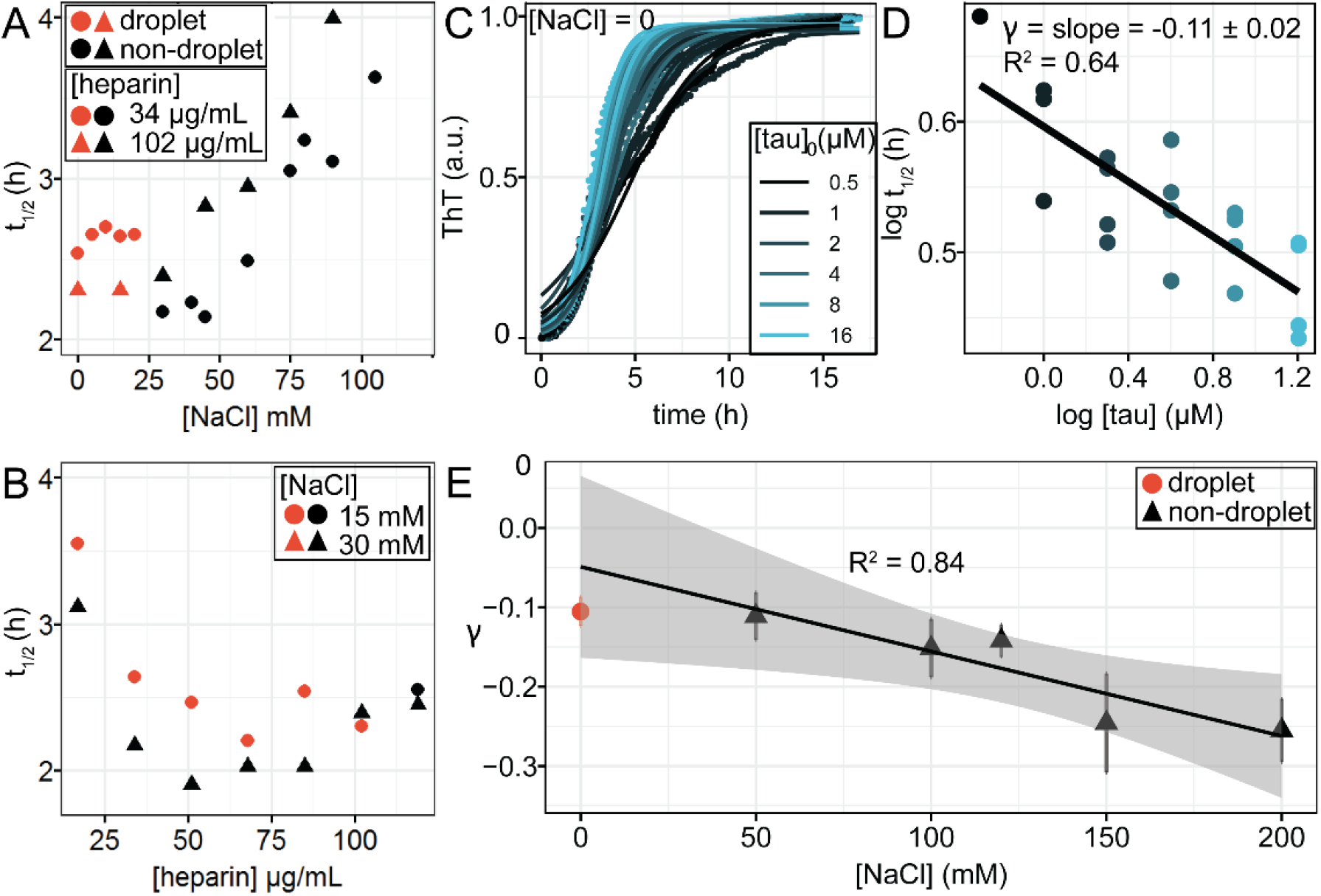
Effects of LLPS on tau-heparin aggregation half time and monomer dependence. **A.** t_1/2_ of tau-heparin aggregation at different [NaCl]. 20 *μ*M tauSS with 34or 102 *μ*g/mL heparin were used. **B.** t_1/2_ of tau-heparin aggregation at different [heparin]. 20 *μ*M tauSS and 15 or 30 mM NaCl were used. **C.** Normalized ThT fluorescence of tauSS-heparin mixture at droplet conditions ([NaCl] = 0) with varying initial tau concentration, [tau]_0_, from 0.5 *μ*M to 16 *μ*M, colored in gradient from black to blue. Readings are marked in points. Solid line shows the fitted results using sigmoid function. Ratio of heparin was fixed at 1.7 *μ*g/mL heparin per 1 *μ*M tau. **D.** log-log plot of t_1/2_ vs [tau]_0_. Data points are t_1/2_’s extracted from fitting (solid lines in A). Solid line is a linear regression of the data points, with R^2^ = 0.64. The slope of log-log plot is denoted scaling exponent γ. **E.** scaling exponent γ of tauSS-heparin at varying [NaCl] conditions. Error bars are standard error of the estimate. Solid line is a linear regression of [NaCl] = 50 mM to 200 mM with R^2^ = 0.84, and shaded area is the confidence interval (α = 0.95) of the regression. In A, B and E, data points in red are conditions with turbidity above 30% maximum and were labeled as droplet conditions.

We point out small but reproducible non-linear dependence of t_1/2_ on [NaCl] under droplet conditions (Figure S5B). Data at non-droplet conditions can be fitted to a linear regression trend line (Figure S5B), which indicates the direct dependence of t_1/2_ on [NaCl]. Comparing with the trend line, aggregation under droplet conditions appears slightly slower than without phase separation.

The aggregation kinetics of many amyloid-forming proteins, including insulin [64], Aβ [65], as well as tau [48], [66], [67], have been modeled by a nucleation-elongation process, in which fibrils form through sequential addition of monomers to the nucleus [68]. The slope of the log-log plot of half time vs initial monomer concentration, denoted scaling exponent *γ*, has been used to identify the dominant microscopic process in the overall reaction [36], [69], [70]. A value for this slope *γ* = 0 implies a completely saturated dominant process [36], [48], meaning the rate of fibril formation is limited by the interaction between the monomer and the interface of the nucleus, and the monomer concentration is so high that the interface is fully occupied, resulting in an apparent independence of t_1/2_ on monomer concentration. Meanwhile, −0.5 < *γ* < 0 implies reduced saturation and a recovery of the monomer dependence [36]. Here, we adopted the nucleation-elongation model for tau aggregation and utilized γ to quantify the effects of LLPS on the monomer dependence of aggregation. We varied the initial monomer concentration of tau, [tau]_0_, keeping heparin at the same ratio R = 1, and recorded t_1/2_. Then, we calculated γ from the slope of the log-log plot of t_1/2_ vs [tau]_0_. We used several [NaCl] values so that aggregation kinetics is monitored under droplet to non-droplet conditions.

We prepared tauSS and heparin samples with [NaCl] = 0. Turbidity readings scaled linearly with [tau]_0_, confirming that droplets formed (Figure S6A). ThT fluorescence of each sample was monitored for 16 hours. The readings were normalized and fitted to extract t_1/2_ (Figure **3**C) (see Materials and Methods). The log-log plot of t_1/2_ vs [tau]_0_ extracted from (Figure **3**C) resulted in data points that can be fit by linear regression with R^2^= 0.64 to extract γ (Figure **3**D). We obtained γ = (−0.11 ± 0.02) at [NaCl] = 0 under LLPS conditions (Figure **3**D). This implies that tau aggregation in LLPS conditions has minimal monomer dependence. We further varied [NaCl] from 50 to 200 mM. At [NaCl] ≥ 50 mM, turbidity readings are independent of [tau]_0_, implying that droplet is absent (Figure S6A). The log-log plot of t_1/2_ vs [tau]_0_ at all [NaCl] conditions from 50 to 200 mM can be fit linearly with R^2^ above 0.64 to obtain γ (Figure S6C). We observed that γ changes from (−0.11 ± 0.03) at [NaCl] = 50 mM to (−0.26 ± 0.04) at 200 mM, and can be linearly fit with R^2^ = 0.84 (Figure 3E). This means the monomer dependence of aggregation at nondroplet conditions increases as the ionic strength increases. Importantly, γ under droplet condition (−0.11 ± 0.02) falls well within the confidence interval of linear regression (Figure **3**E), implying γ is altered by the change in ionic strength, but not by the state of LLPS itself. In other words, the monomer dependence of the microscopic process depends on the inter-molecular electrostatic interaction strength modulated by ionic strength, but the increased protein concentration under LLPS is insufficient in and of itself to modulate the microscopic processes governing tau aggregation.

### Aggregation-signature conformation is independent of LLPS

We have previously identified a structural signature of aggregation-prone tau conformers, which consists of an opening of the flanking regions of the hydrophobic PHF6 and PHF6* hexapeptide segments of tau [37]. We here assessed whether a similar conformational change is triggered under LLPS conditions. Double electron-electron resonance (DEER) spectroscopy is a powerful tool that measures intramolecular distances to report on a local conformational ensemble [38], [39]. Here we used DEER to measure the distribution of intra-molecular distances of the tau ensemble between residues 300 and 313, flanking both sides of the PHF6 segment (i.e. ^306^VQIVYK^311^). We referred to the construct containing spin labels at site 300 and 313 as tauSL_2_. The concept of this experiment was to measure conformation around tau’s PHF6 segment immediately after adding a cofactor that either triggers aggregation, LLPS, or both. The conformation of the PHF6 segment is used as a proxy to evaluate aggregation-prone conformations of tau, as described in previous studies [37]. For reference, two control samples were introduced: tauSL_2_ in the absence of polyanion was used to represent PHF6 conformation of tau monomer in solution (referred to as monomer); tauSL_2_ in fully aggregated tau-heparin fibrils (24h incubation) was used to represent tau conformation in fibrils. The goal of this set of experiment was to reveal whether the soluble aggregation-prone conformation can be correlated with LLPS.

The first DEER sample (sample I) was prepared by mixing the tauSL_2_ with heparin at a ratio R = 1. A clear solution immediately became turbid, and light microscope confirmed the abundance of droplets (Figure 4A, column 1). Replica were prepared and incubated with ThT. Fluorescence showed that amyloid aggregates became gradually abundant after overnight incubation. Within 5-10 minutes of preparation, the sample was flash frozen for DEER measurements, while the ThT signal was minimal revealing the absence of significant quantities of amyloid aggregates (Figure 4B, column 1). Both the DEER time trace (Figure 4C, column 1) and the extracted distance distribution, P(r) (Figure 4D, column 1) showed extended conformations, similar to conformation observed in fully aggregated sample.

**Figure 4.**
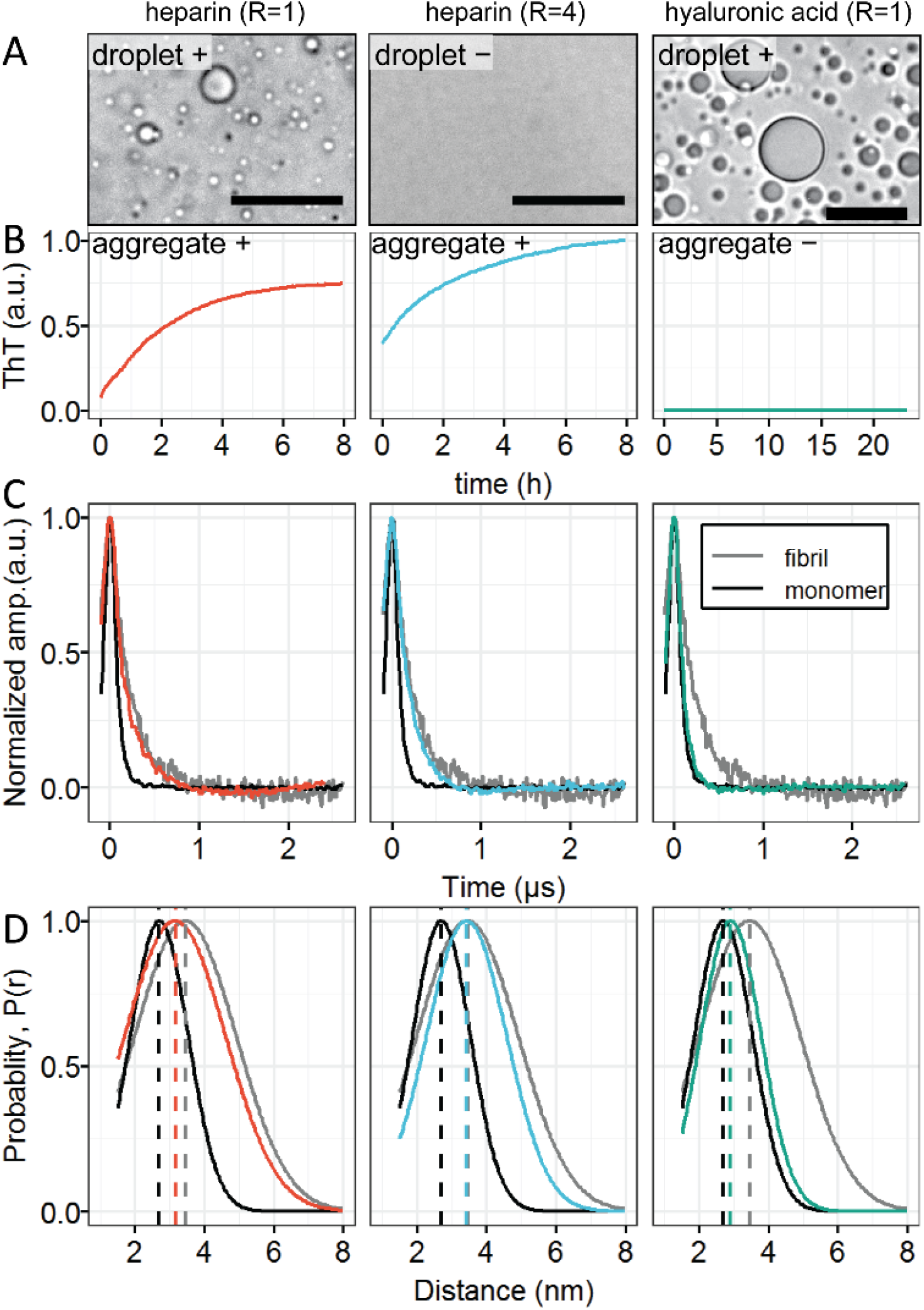
LLPS dependence of tau aggregation-signature conformation. **A.** Bright field microscope images of tau-heparin sample at R = 1, R = 4 and tau-hyaluronic acid sample at R = 1 in the DEER conditions. [tau]=220 μM. Scale bars are 25 μm long. **B.** ThT fluorescence of samples in A during incubation. **C.** DEER time domain signal. **D.** Corresponding distance distribution *P*(*r*).

This observation is consistent with the previous report that tau adopts aggregation-prone conformation at early stages of amyloid aggregation. The question is whether droplet formation is related to, or presumably facilitating such conformation change. We next prepared the second DEER sample by introducing 4 times more heparin (sample II) to reach a ratio R = 4. Under this condition, LLPS did not occur (Figure 4A, column 2), while ThT assay shows that aggregation is still triggered (Figure 4B, column 2). Despite the absence of droplet, DEER signal and P(r) of the sample II resembled those of fibril sample (Figure 4C, D, column 2). Clearly, droplet formation is not required for tau to adopt the aggregation-prone signature conformation.

Sample III was prepared by incubating tau with hyaluronic acid at charge ratio R = 1. Hyaluronic acid shares the same backbone and similar molecular weight as heparin (used in sample I and II), but is free of sulphate groups (Table 2). By doing so, we created LLPS (Figure 4A, column 3) without triggering amyloid aggregation (Figure 4B, column 3). Despite the presence of droplet, the DEER signal and P(r) were found to be similar to those of the monomer sample (Figure 4C, D, column 3). Clearly, droplet formation itself is not sufficient to drive tau aggregation-prone conformation change. Together, these results show that LLPS does not directly result in tau conformational changes associated with enhanced aggregation propensity.

### Amyloid aggregation can be modulated independently of droplet formation

Knowing that LLPS has no direct impact on the conformational state of tau, aggregation reaction rate of tau or final extent of amyloid aggregation, we hypothesized that LLPS and amyloid aggregation are driven by distinct interactions. To test this hypothesis, we modulated tau amyloid aggregation without disturbing the charge-charge interactions between tau and polyanion. We then measured and compared their LLPS and amyloid aggregation under the same experimental conditions.

We first compared 6-O-desulfated, 2-O-desulfated and N-desulfated-reN-acetylated heparins, named 6OD, 2OD and ND, respectively (Table 2). They all have glycosaminoglycan backbone with 2 sulfate and 1 carboxyl charged groups, resulting in an identical charge density (3 e^−^ per disaccharide), but with different charge location on the backbone (Table 2). According to the supplier, these desulfated heparins are products of the same heparin after reaction with different enzymes. Consequently, they share similar molecular weights. We identify the charge ratio of heparin to tau, R, to represent its stoichiometry. When mixed with tauS under charge neutrality conditions, i.e., R = 1, all three heparin variants make droplets with tau (Figure 5A). Furthermore, the titration of different ratio R showed that desulfated heparins have indistinguishable turbidity on its own for all R in the interval of 0 to 2 (Figure 5B). In other words, these three desulfated heparins have similar LLPS propensity. However, incubation of tauS overnight with the desulfated heparins at R = 1 resulted in dramatically different ThT fluorescence (Figure 5C). Notice 6-O-desulfated heparin has been reported to have lowest affinity with full length tau 2N4R, compared with 2-O-desulfated and N-desulfated-reN-acetylated heparin [40], while in our experimental conditions 6-O-desulfated heparin showed highest aggregation propensity (Figure 5C). Regardless of the ordering, the observation is clear that the different desulfated heparins yield similar turbidity upon LLPS with tau but different ThT fluorescence, demonstrating that tau fibrillization can be modulated by arranging the charged sufate group position on the glycosaminoglycan backbone, without interfering with and without being affected by, LLPS.

**Figure 5.**
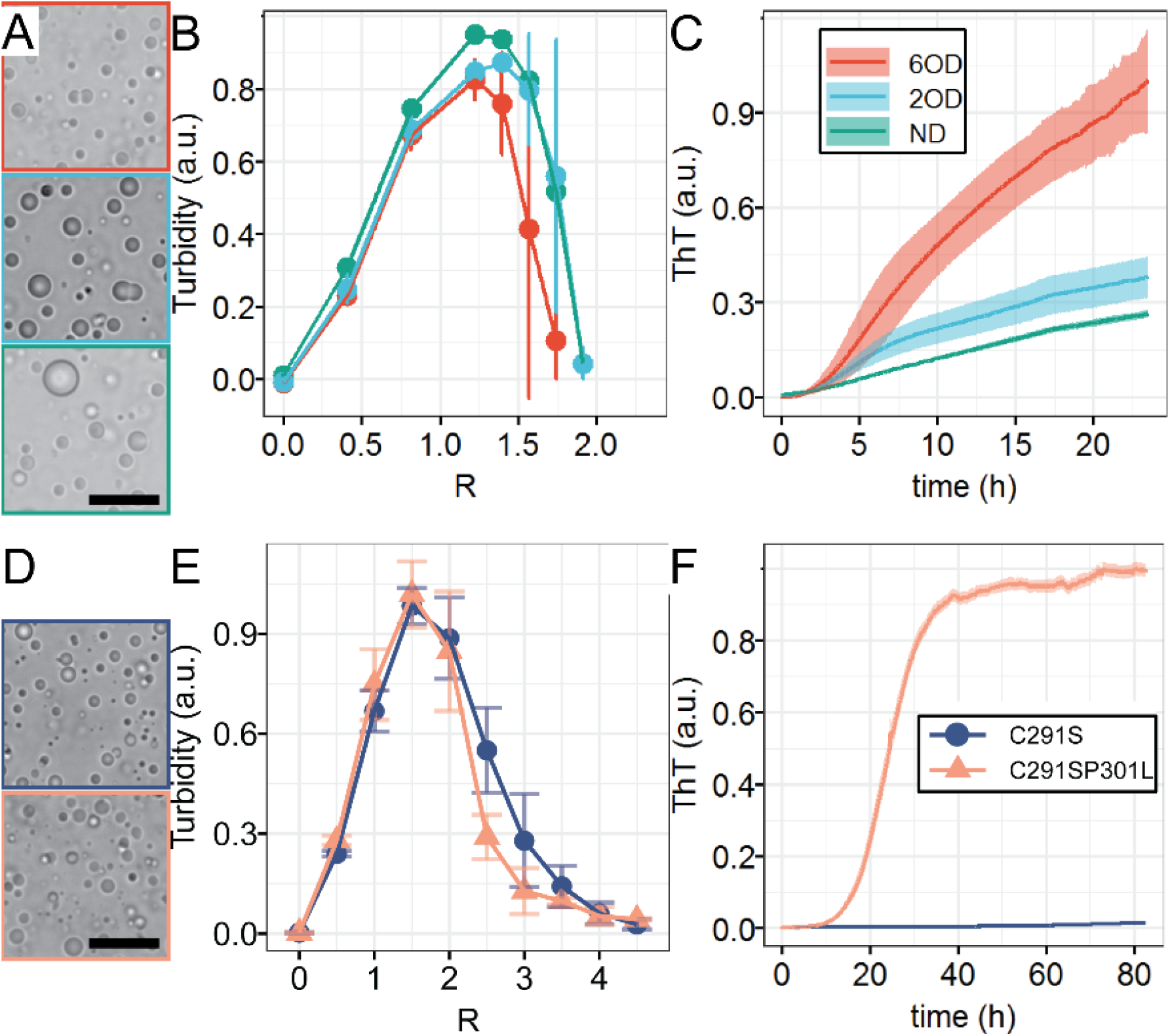
Dependence of LLPS on modulating amyloid aggregation. **A.** Representative bright field microscope images of tau-polyanions. 6OD, 2OD, ND are 6-O-desulfated heparin, 2-O-desulfated heparin and N-desulfated reN-acetylated heparin, respectively. Desulfated heparins were derived from heparin of same molecular weight. [tau] = 100 μM, [NaCl] = 0, R = 1. **B.** charge ratio, R, dependence of turbidity for tauS titrated with different desulfated heparins. [tau] = 20 *μ*M, [NaCl] = 0. **C.** ThT fluorescence of 20 μM tauS incubated with different desulfated heparins over 24 hours. R = 1, [NaCl] = 0. **D.** Representative bright field images of tau-poly(U) RNA for two tau variants. **E.**Turbidity of tau-poly(U) RNA with different tau variants at varying R. **F.** ThT fluorescence of tauRNA for different tau variants. Scale bar lengths in both A and B are 20 *μm*. Error bars in both B and E show standard deviation of 3 independent repeats. Shaded areas in both C and F show standard deviation of 3 replicas.

We then asked whether the disease-associated P301L tau mutation influences the propensity for tau to form LLPS. We compared RNA LLPS with tauS vs. with tauSP301L. TauSP301L differs from tauS by a single P301L mutation—a well-known disease mutation found in fronto temporal dementia [41], but has unaltered net positive charge (Table 1). When mixed with RNA at ratio R = 1, both tauS and tauSP301L form droplets (Figure 5D). Further titrating tau with varying R showed that both tau variants have similar LLPS propensity (Figure 5E). However, incubation of two tau-RNA sample at R = 1 showed that tauSP301L-RNA has ThT readings by 2 orders of magnitude higher than tauS-RNA (Figure 5F). The much higher aggregation propensity of P301L variants of tau is well known. These results together proved that amyloid aggregation can be modulated by changing either the polyanion chemistry or the tau protein mutation without affecting LLPS properties.

### LLPS does not directly promote seeding

Tau amyloid aggregation can be seeded, by adding premade fibrils in the aggregation reaction, both *in vivo* [42]–[45] and *in vitro* [46]–[48]. *In vitro* seeding has a low efficiency that was shown to be increased in the presence of a mild cofactor such as RNA [46], [49]. We asked the question whether LLPS is involved and facilitatory for such enhancements. To solve this question, we compared seeding enhancement with RNA between droplet and non-droplet conditions. TauSSP301L was incubated with heparin overnight to prepare fully aggregated tau fibrils as the seed. 5% seed was incubated with tauSSP301L to trigger seeding reaction with and without RNA. This reaction was carried out at several [NaCl], two of which conditions produced droplets (0 and 50 mM, Figure S7) while two other concentrations did not (100 and 150 mM). As expected, ThT fluorescence of seeded aggregation with RNA as an added cofactor was significantly enhanced compared to seeding without RNA (Figure 6). This enhancement, although decreasing with increasing salt, was still visible in non-droplet conditions and therefore did not appear to be correlated with the formation of LLPS. These results demonstrated that LLPS is not necessary for cofactor-enhanced *in vitro* seeding.

**Figure 6.**
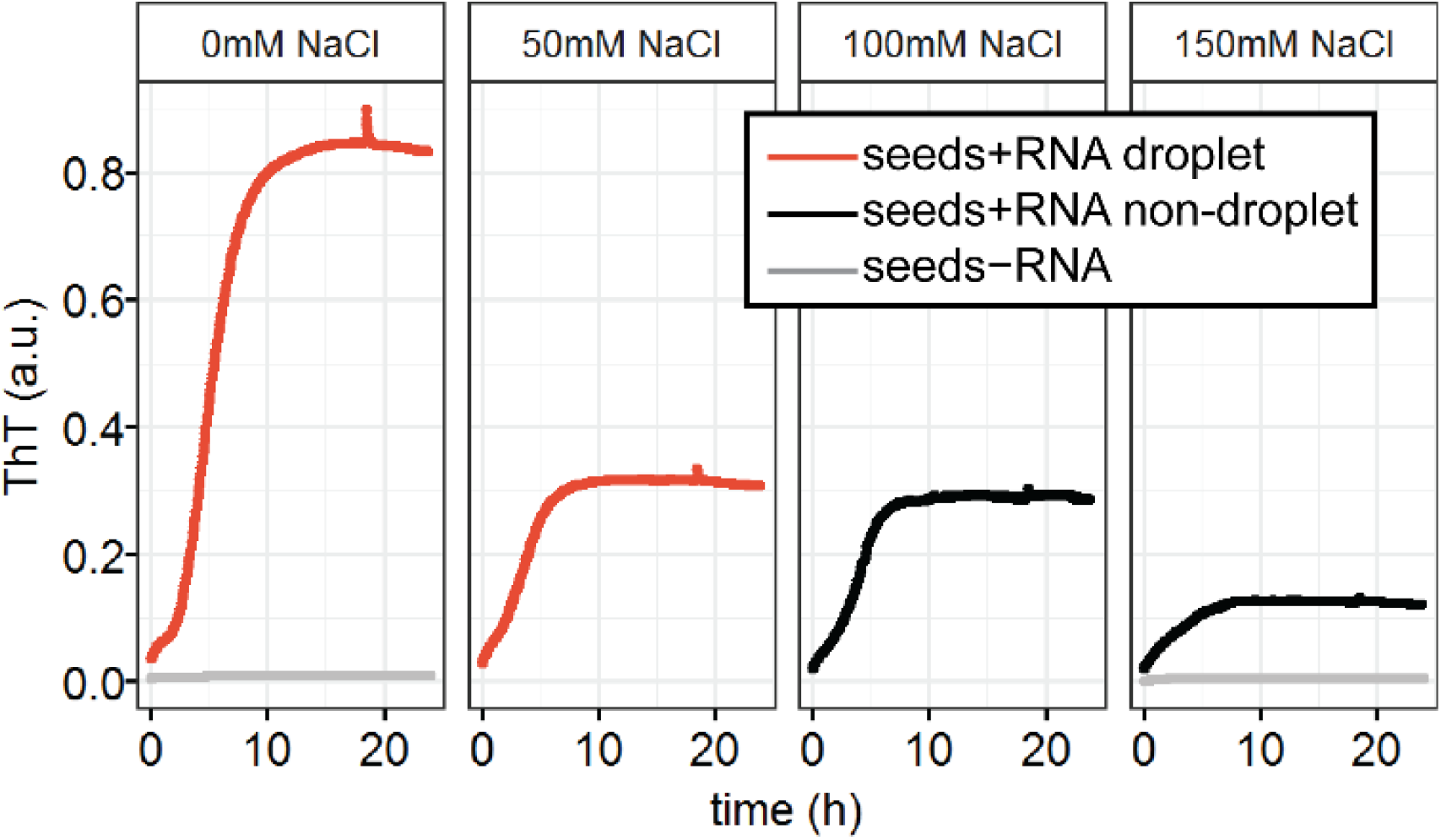
LLPS dependence of seeding. 20 *μ*M tauSSP301L was incubated overnight with pre-formed seed (5% molar) at various [NaCl] in the presence (red and black lines) or not (grey lines) of 60 *μ*g/mL poly(U) RNA while measuring ThT fluorescence. Droplet conditions were assigned to samples whose normalized turbidity was above 0.3 (0 mM and 50 mM NaCl), and were colored in red, while non-droplet conditions were colored in black (100 mM and 150 mM NaCl). Seeding enhancement was observed with addition of RNA, both in and out of droplet conditions, showing that this enhancement is not due to LLPS.

## Discussion

Tau droplets and amyloid aggregates have been found to occur simultaneously [28], [29], but it was unclear whether droplets are on pathway towards aggregates. Our work demonstrated that LLPS and amyloid aggregation can be two independent processes that influence each other indirectly. We proposed possible underlying mechanisms.

First, we propose that LLPS and tau amyloid aggregation are driven by distinct interactions. This is supported by the observation that tau-heparin amyloid aggregation occurs at ionic strength up to 160 mM, while [NaCl] = 50 mM is enough to completely eliminate LLPS (Figure 2). The common preference for low ionic strength is consistent with previous studies [50], [51], and explains why these two processes are often observed under overlapping experimental conditions. However, the Debye lengths (the characteristic distance between two point charges in solution beyond which electrostatic interaction becomes negligible [52]) of 160 mM and 50 mM NaCl are approximately 0.76 nm and 1.36 nm [52], corresponding to the estimated contour length of 2 and 3.6 amino acids, respectively [53]. Therefore, it is likely that amyloid aggregation is mediated by a shorter range of electrostatic interactions than LLPS conditions can invoke or modulate. Furthermore, the stability of amyloid aggregate is known to originate from the inter-backbone hydrogen bond network [54], [55] assisted by specific side-chain interactions [56], [57]. In contrast, complex coacervation of tau and polyanion is dominantly controlled by long-range, weaker and multivalent electrostatic interactions [30], [31]. This explains the result that tau amyloid aggregation is sensitive to change in the site-specific charge location or distribution on polyanion cofactors, and is sensitive to specific mutation on tau (Figure 5), while LLPS is more tolerant to those variations.

Second, tau-polyanion LLPS and amyloid aggregation may have separated pathways. Aggregation-prone conformation is associated with amyloid aggregation regardless of LLPS (Figure 4), suggesting LLPS does not directly change tau conformation. This is consistent with the recent reports that the conformations of polyelectrolyte chains under complex coacervate is highly similar to that in solution [58], [59]. Meanwhile we observed that the optimal stoichiometry of heparin that facilitates amyloid aggregation is very different from that for LLPS (Figure 2). This is consistent with the previous observations that tau-polyanion LLPS favors stoichiometry that meets net charge balancing [30], while tau-heparin amyloid aggregation proceeds via a defined molar ratio of tau and heparin, likely mediated by directed conformational templating [49], [50]. The distinct conformation change and optimal stoichiometry suggest these two processes are governed by distinct mechanisms.

Notice that the independence of LLPS on amyloid aggregation might be specific to the tau-cofactor system that proceed via complex coacervation. LLPS can be achieved using solely tau with specific truncation [28], crowding reagent [60] or post-translational modifications (PTMs) [29]. Such simple coacervation relies on tau-tau interactions that are unlikely dominant in complex coacervation. This explains the conformation changes [28], [34] and subsequent amyloid aggregates [28], [29] observed in the simple coacervation of tau.

While our work shows that LLPS and aggregation have distinct driving forces, the two processes most likely interfere when occurring simultaneously. We observed a small but significant change in the aggregation half time within droplets (Figure S5). We proposed such change may result from two counter-acting properties of droplets: 1) LLPS creates a high concentration environment, which increase collision probabilities and may speed up amyloid aggregation; 2) Such compartment may also have high viscosity, which reduces the diffusion of tau, and therefore slows down aggregation. The potential opposite effects of these two properties explain the small change in the reaction rate for aggregation in LLPS.

Complex coacervation permits the concentration and colocalization of two or more compounds that have matching charge patterns into so-called membrane less organelles [13]. This concentration effect is likely to have a much more prominent effect in a cellular context than under *in vitro* experiment where only two purified compounds are present (tau and RNA) at 10s of μM. Firstly, the concentration of unbound tau in neurons can be heterogeneous and orders of magnitude lower than values in our experiments, giving little probability for tau-tau collisions and interactions (necessary step of aggregation) to occur, unless there is a mechanism of colocalization of tau, possibly in the form of LLPS. Secondly, it has been recently pointed out that tau fiber might naturally incorporate cofactors that may change the dependence between the aggregation and LLPS processes [10], [49]. In addition, the colocalization and concentration of enzymes could promote PTMs, such as phosphorylation [61], that can be connected to tau aggregation. LLPS might be an essential step to mix tau and pathologically relevant cofactors that become part of or facilitate tau aggregation, or regulate other essential processes. What our study is showing is that the physical properties of LLPS of tau is not in and of itself providing a mechanistic basis and driving factors for enhanced aggregation kinetics or propensity. If there is a role that LLPS plays in the aggregation process of tau, it would be on the basis of the interplay of biological factors.

## Conclusion

In this work we showed that tau is able to undergo LLPS with a wide range of polyanions tested. Among these polyanions, only a subset can facilitate tau amyloid aggregation. Under LLPS conditions, amyloid aggregates do colocalize with the tau droplets. By establishing a 2-dimensional map (ionic strength and stoichiometry) of LLPS and amyloid aggregation, we found that the state of LLPS does not change the extent of amyloid aggregation. We found that the half time for aggregation kinetics is insignificantly altered, nor is the monomer dependency of the dominant microscopic processes during aggregation affected in the LLPS state. By double electron-electron resonance spectroscopy, we showed that LLPS does not induce the characteristic tau conformational rearrangements consisting in an opening of the PHF6(*) segments, which have been previously demonstrated to be aggregation-prone signatures for tau. We also modulated amyloid aggregation propensity by either changing the polyanion type or by introducing disease mutations to tau without affecting its length or charge density. By doing so we found that the state of LLPS does not affect the kinetics nor propensity for amyloid aggregation. We furthermore studied the seeding activity of tau under aggregation-prone conditions, and found that LLPS also does not directly enhance seeding activity.

## Acknowledgement

The authors acknowledge support for the studies of tau aggregation and LLPS mechanisms from the National Institutes of Health (NIH) (grants R01AG056058(S.H.)) and the Tau consortium (http://www.tauconsortium.org) from the Rainwater foundation.

## Materials and Methods

### Protein expression and purification

In this work, we used an N-terminal truncated form of tau, tau187, containing the microtubule binding domain (residues 255-441 from the longest human tau isoform 2N4R with a 6× His-tag at the N-terminus). Site-directed mutagenesis was applied to prepare the following mutated variants of tau187: tauS contains C291S mutation; tauSP301L contains C291S and P301L mutations; tauSS contains C291S and C322S mutations; tauSSP301L contains C291S, C322S and P301L mutations; tau187V300CV313C contains C291S, C322S, V300C and V313C mutations. The names and estimated charge properties are listed in (Table 1).

We followed the previously reported methods for expression and purification of tau187 [35], [49], [62], [63]. DNA variants were constructed and used to transfect E. coli BL21 (DE3) cells. Cells were stored as frozen glycerol stock at −80 °C. Cells from glycerol stock were grown in 10 mL luria broth (LB, Sigma Aldrich, L3022) overnight and then used to inoculate 1 L of fresh LB. Cells were grown at 37 °C, 200 rpm with addition of 10 μg/mL kanamycin (Fisher Scientific, BP906) until optical density OD_600_ reached 0.6–0.8. Next 1 mM isopropyl-ß-D-thiogalactoside (Fisher Bioreagents, BP175510) was added to induce expression. After incubation for 2–3 h, cells were palleted with centrifugation at 5000 g at 4 °C for 20 min. Harvested cells were resuspended in lysis buffer (Tris-HCl, pH = 7.4, 100 mM NaCl, 0.5 mM DTT, 0.1 mM EDTA) added with 1 Pierce protease inhibitor tablet (Thermo Scientific, A32965), 1 mM PMSF, 2 mg/mL lysozyme, 20 μg/mL DNase (Sigma, DN25) and 10 mM MgCl_2_ (10 mM). Samples were incubated on ice for 30 min, then frozen in liquid nitrogen and thawed in room temperature for 3 cycles. Cell debris was removed by centrifugation at 10,000 rpm at 4 °C for 10 min. 1 mM PMSF was added again and samples were heated at 65 °C for 12 min and cooled on ice for 20 min. Cooled samples were then centrifuged at 10,000 rpm for 10 min to remove the precipitant. The resulting supernatant was incubated overnight with Ni-NTA resins pre-equilibrated in buffer A (20 mM sodium phosphate, pH = 7.0, 500 mM NaCl, 10 mM imidazole, 100 μM EDTA). The resins were loaded to a column and washed with 20 mL of buffer A, 25 mL buffer B (20 mM sodium phosphate, pH = 7.0, 1 M NaCl, 20 mM imidazole, 0.5 mM DTT, 100 μM EDTA). Purified protein was eluted with 15 mL of buffer C (20 mM sodium phosphate, pH = 7.0, 0.5 mM DTT, 100 mM NaCl, 300 mM imidazole). Eluents were analyzed by SDS-PAGE to collect the pure fractions. Proteins were then buffer exchanged into DTT-free working buffer (20 mM HEPES, pH 7.0) and immediately frozen and kept at −80 °C until usage.

### Preparation of tau-polyanion droplets and microscope imaging

We used the following polyanions in this work for studying LLPS and amyloid aggregation: PolyA RNA (MW 66 ~ 660 kDa, Sigma, P9403); PolyU RNA (MW 800~1000 kDa, Sigma, P9528); PolydA single strand DNA (MW 83~165 kDa, Sigma, 10223581001); Hyaluronic acid (MW 8~15 kDa, Sigma, 40583). Besides, heparin, 6-Odesulfated heparin, 2-O-desulfated heparin and N-desulfated reN-acetylated heparin are all polydisperse, with average MW ~15 kDa and from Galen laboratory supplies under the reference number, HEP001, DSH002/6, DSH001/2 and DSH004/NAc, respectively. The primary chemical structure and charge properties were listed in (Table 2).

We followed the previously reported method for preparing tau-polyanion droplets for microscope imaging [30], [31]. The charge per mass at neutral pH of all tau variants was estimated using Innovagen’s Peptide Property Calculator (http://pepcalc.com/). The charge per mass for polyanions were estimated assuming all charged groups are fully ionized and negatively charged. For microscope imaging, polyanions were mixed with tau in certain mass ratio, as stated. Images were taken within 10 minutes after mixing. Bright-field images were acquired using an inverted compound microscope (Olympus IX70). Confocal images were acquired using a spectral confocal microscope (Olympus Fluoview 1000). Images of ThT fluorescence were taken using λ_emission_=485 nm.

### Turbidity measurement and ThT assays

Tau, polyanions and NaCl were mixed and pipetted onto a micro plate (Corning, 3844). Absorbance at 500 nm and fluorescence intensity (excitation 440 nm, emission 485 nm) were both monitored using a Bio-Tek Synergy 2 microplate reader, with temperature maintained at 26.0 °C. Absorbance was used as turbidity reading, while fluorescence used as ThT fluorescence.

Along with each run of microplate reader, an equal volume of buffer with same concentration of ThT was added to the plate and used as control. Turbidity reading of the control were subtracted from the turbidity reading of each sample. ThT fluorescence reading from each sample were corrected by the control reading, in order to reduce the artifact from lamp drift/fluctuation, following the two equations below.

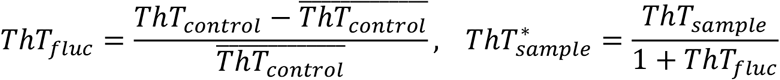

where ThT_control_ is the fluorescence reading of the control samples; 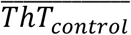 is the averaged value throughout the entire course of measurement; ThT_fluc_ is the drift/fluctuation of the lamp; ThT_sample_ and ThT^*^_sample_ are fluorescence reading of the sample before and after correction, respectively.

We applied the following steps to extract the half time and maximum ThT fluorescence. First, for each well of the microplate, the first 10 minutes of the above lamp-corrected readings were averaged and used as a baseline, assuming no amyloid aggregates formed during this time. Next the baselines of each well were subtracted from the ThT readings.

Third, the maximal ThT readings of the entire microplate during the entire incubation was used to divide all the readings, scaling to 0-1. Fourth, the following sigmoid function was used to fit all the readings, using Levenberg-Marquardt algorithm, provided by nls.lm function in minpack.lm package, with initial guess (*A* = 1, *t*_1/2_ = 5, *k* = 1)

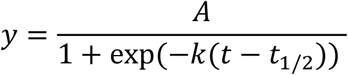

where the fitted A and t_1/2_ were used as maximum ThT fluorescence and half time, respectively. The fitted results were visually examined to ensure a good fit.

### Double Electron Electron Resonance (DEER)

Tau187V300CV313C was expressed as other tau187 mutants and spin labeled as follow. Tau was freshly purified and treated with 5 mM TCEP. TCEP was removed using a PD-10 desalting column. Immediately after PD-10, 10× to 15× molar excess of MTSL ((1-Acetoxy-2,2,5,5-tetramethyl-δ-3-pyrroline-3-methyl) Methanethiosulfonate, Toronto Research Chemicals, O875000) to free cysteine was incubated with the protein at 4C overnight. After incubation, excess MTSL was removed using a PD-10 desalting column. Labelling efficiency, defined as the molar ratio of tethered spin labels over the cysteines, was measured to be 50-60%Tau187C291SC322S (cysteine-less tau construct, referred to as cysless). Both two protein stocks were concentrated, and buffer exchanged against D_2_O-based buffer (20 mM HEPES in D_2_O) using Amicon centrifugal concentrators (10 kDa cutoff). A 1:22 molar ratio of tau-SL_2_:tau-cysless sample of 10 μM tau-SL_2_ and 2100 μM tau-cysless was mixed with polyanions at designated concentrations and incubated for designated time. 28 μL samples were mixed with 12 μL D_8_-glycerol (Sigma-Aldrich) before transferring to a quartz tube (2 mm i.d.) and frozen using liquid nitrogen.

Four-pulse DEER experiments were carried out at 85 K using the Q-band Bruker E580 Elexsys pulse EPR spectrometer operating at ~34 GHz and equipped with a 300 W TWT amplifier. The following DEER pulse sequence was used: π_obs_/2 − τ_1_− π_obs_ − (t-π_pump_) − (τ_2_-t) − π_obs_ − τ_2_ − echo. Rectangular observe pulses were used with lengths set to π_obs_/2 = 10-12 ns and π_obs_= 20-24 ns. A chirp π_pump_ pulse was applied with a length of 20-24 ns and a frequency width of 60 MHz. The observe frequency was 90 MHz higher than the center of the pump frequency range. τ_1_ was 180 ns and τ_2_ was set to 2.4 ms. The DEER experiment was accumulated for ~12 h. The background-subtracted data were fitted assuming a Gaussian distribution of the inter-spin distances. The analysis was done using the LongDistance software (http://www.chemistry.ucla.edu/directory/hubbell-wayne-l).

### In vitro seeding

20 μM tauSSP301L was preincubated with 5 μM heparin overnight to prepare tau-heparin fibrils. Pre-incubated tau-heparin fibrils were used as the seed for seeding experiment. 5% (molar ratio) of seed was mixed with freshly prepared tau monomer and designated concentration of polyanions in the presence of 20 μM ThT and different concentration of NaCl. Both turbidity and fluorescence were monitored using microplate reader.

---

**Figure S1.**
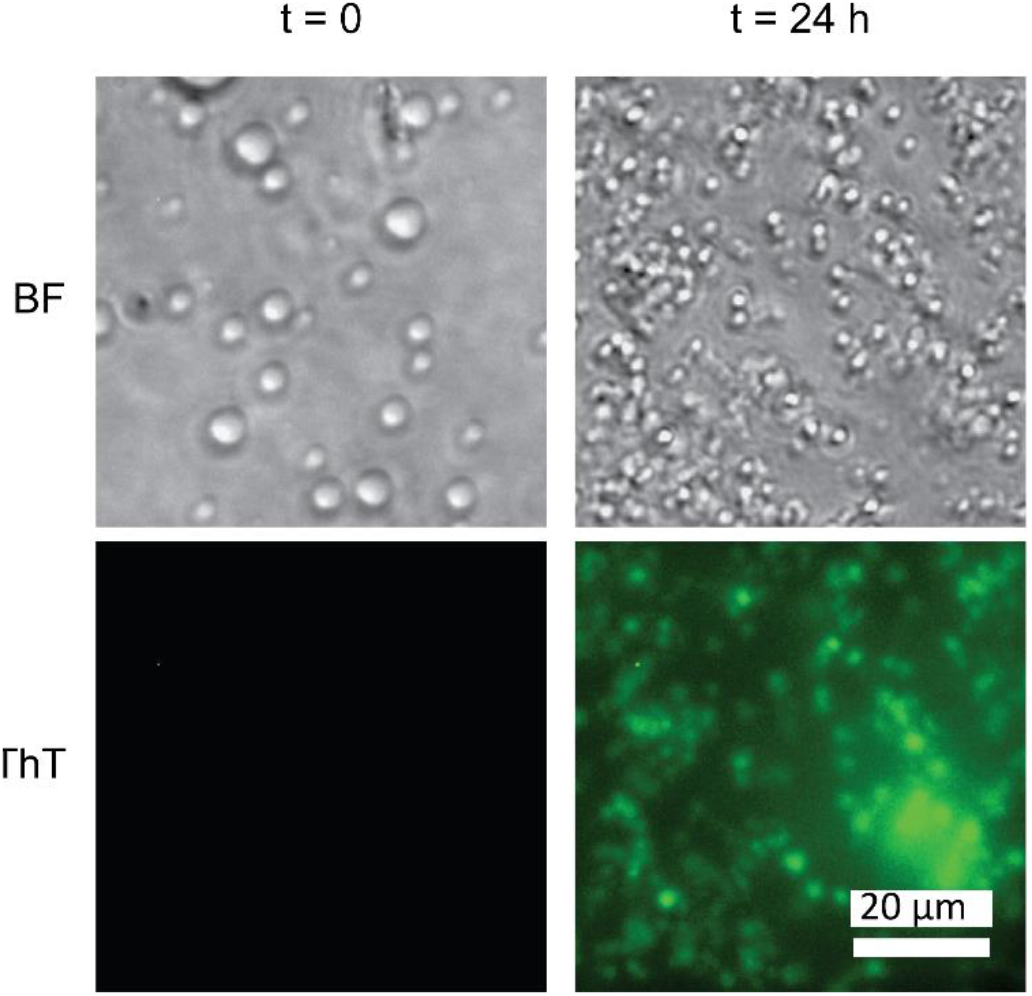
Microscope images of tau-heparin LLPS and amyloid aggregation at room temperature. 100 μM tauS was incubated with 170 μg/mL heparin and 10 μM ThT at room temperature for 24 h. Bright field (BF) images and fluorescence of ThT (λ_emission_=485 nm) were taken immediately after mixing (t = 0h), and 24 hours after incubation (t = 24 h). Scale bar is 20 μm long.

**Figure S2.**
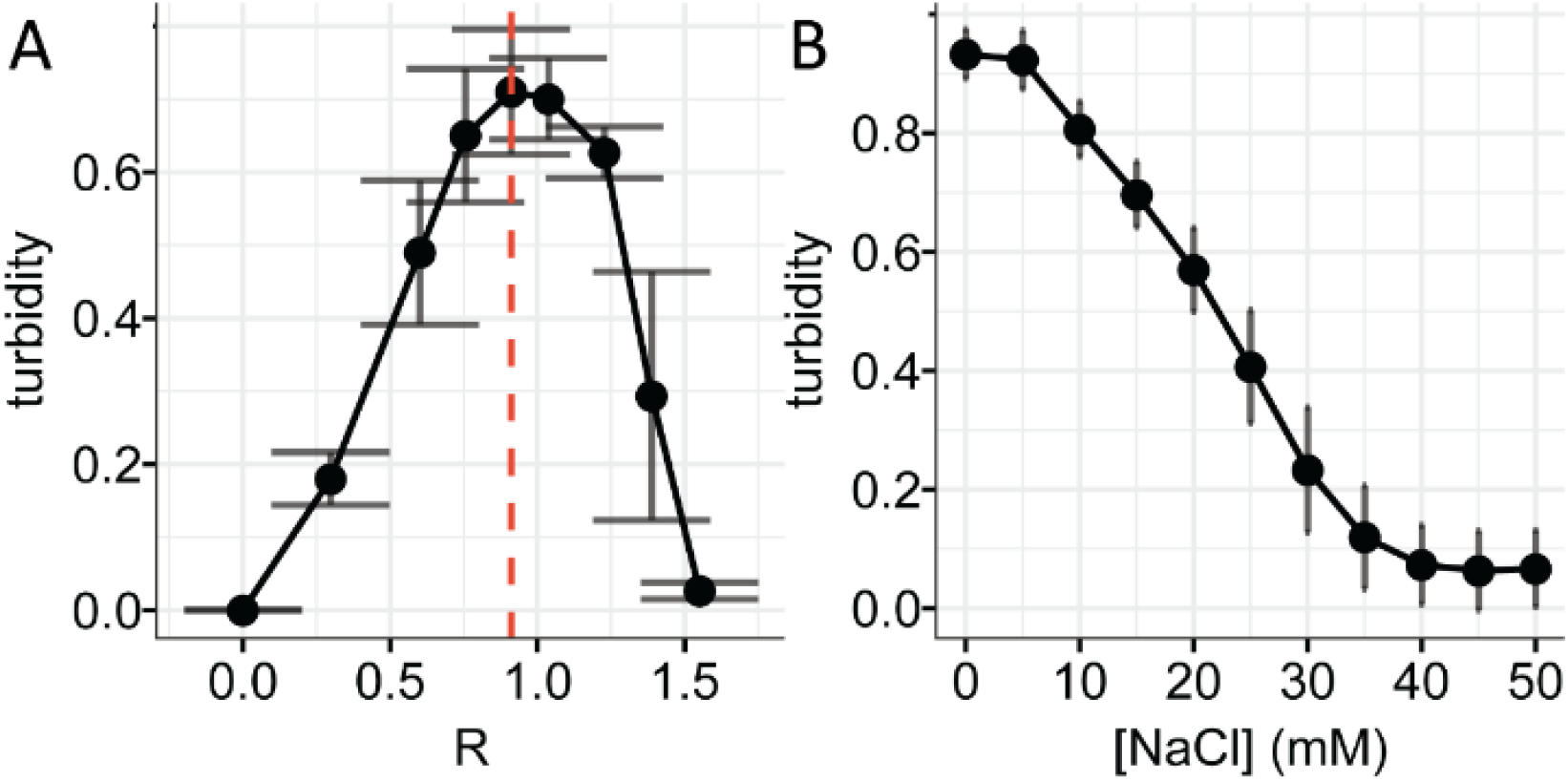
Turbidity dependence of tau-heparin LLPS at varying charge ratio and ionic strength. **A.** Turbidity of tauS-heparin mixture at varying charge ratio (R). [tau] = 20 μM, [NaCl] = 0. R = 1 corresponds to 1.7 μg/mL heparin per 1 μM tau. Dashed line shows the position of the peak reading. **B.** turbidity of tau-heparin at varying [NaCl] from 0 to 50 mM. [tau] = 20 μM, R = 1. Error bars are standard deviation of 3 independent repeats.

**Figure S3.**
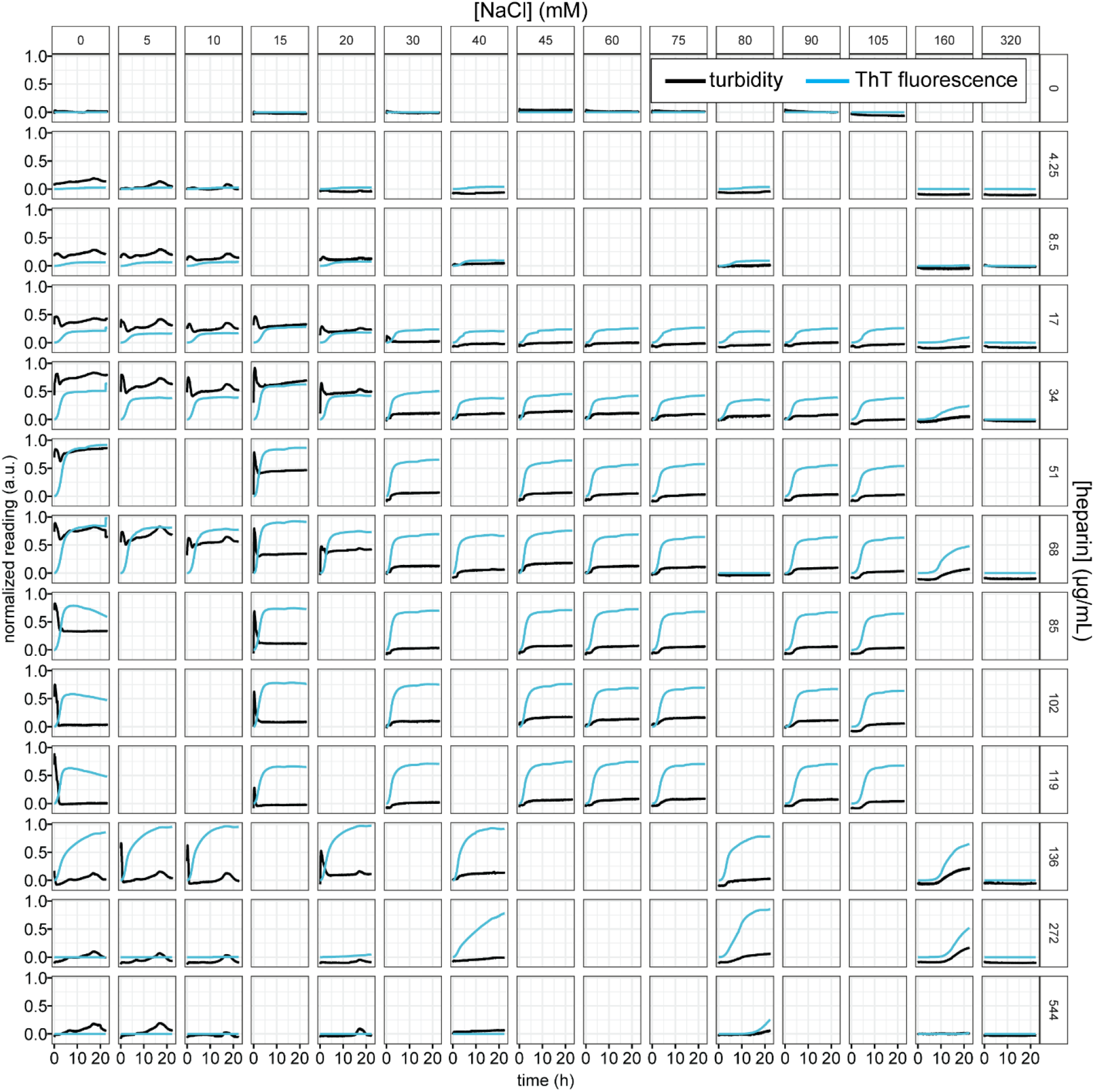
ThT fluorescence and turbidity data of Figure 2. 20 μM of tauSS was incubated with heparin at varying [heparin] and [NaCl]. Turbidity (black) and ThT fluorescence (blue) of each sample were monitored and scaled to 0~1. Different rows correspond to varying [heparin] from 0 to 544 μg/mL. Different columns correspond to varying [NaCl] from 0 to 320 mM.

**Figure S4.**
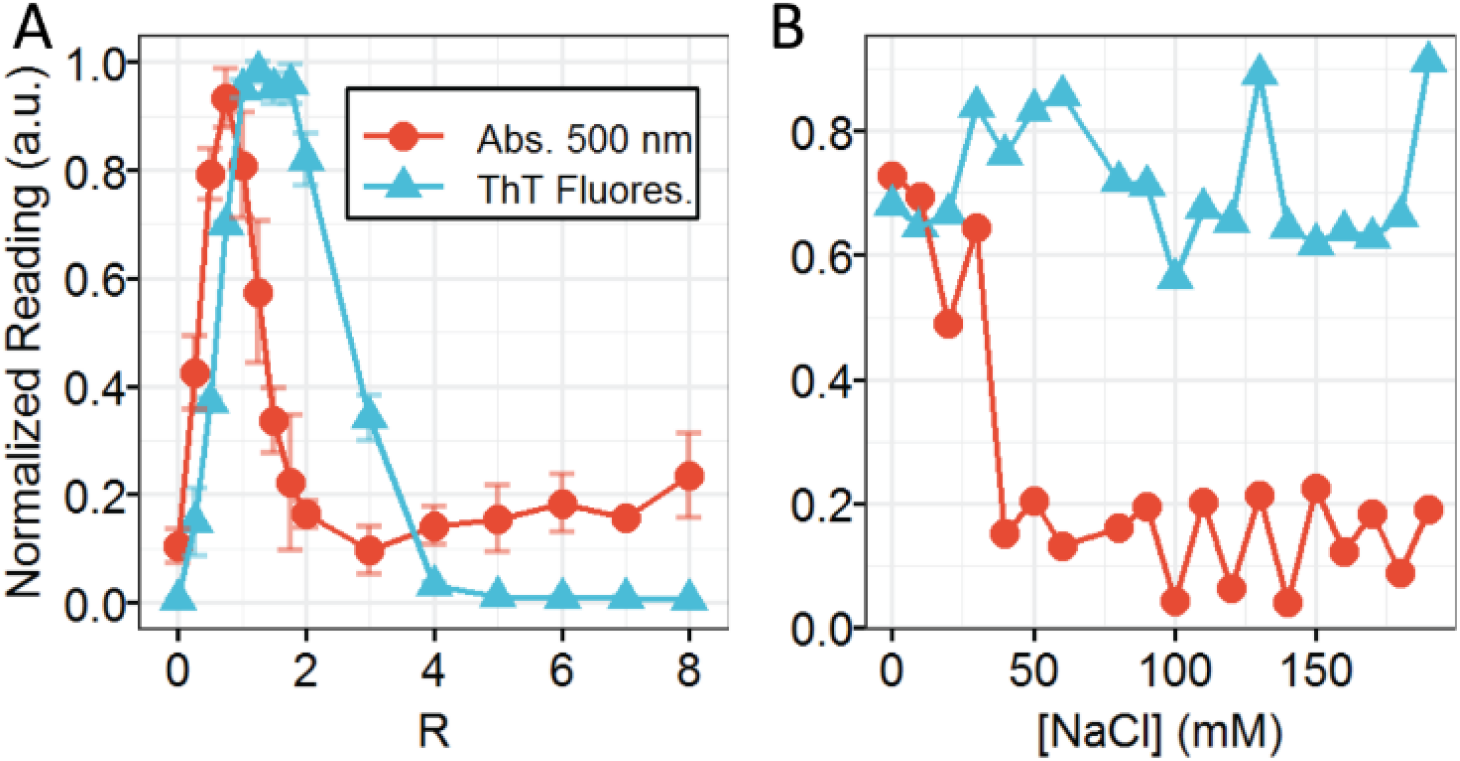
Turbidity and ThT fluorescence of tauS-heparin. **A**. Normalized Turbidity and ThT fluorescence of tau187C291S-heparin aggregation at different charge ratio, R. [tau] = 20 μM, [NaCl] = 0 mM. R = 1 corresponds to [heparin] = 34 μg/mL. Error bars show standard deviation of 3 replicates. **B.** Normalized turbidity and ThT fluorescence of tau-heparin aggregation at different [NaCl]. [tau] = 20 μM. Charge ratio R = 1.

**Figure S5.**
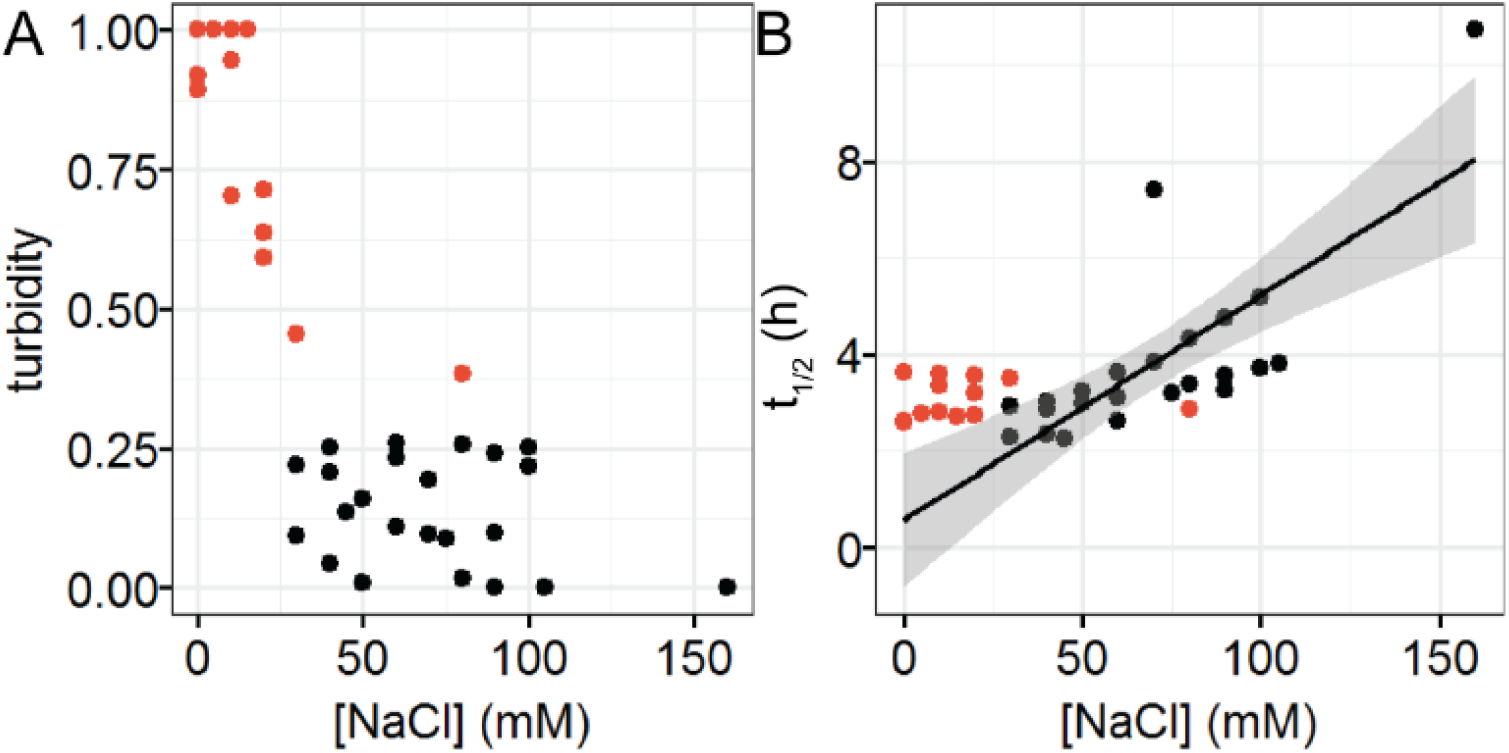
Effects of LLPS on tau-heparin aggregation half time. **A.** Normalized turbidity reading of tauSS-heparin mixture at varying [NaCl]. Data points are independent repeats. **B.** t_1/2_ of tauSS-heparin aggregation at different [NaCl]. 20 *μ*M tauSS with 34 *μ*g/mL heparin were used. In both A and B, data points with normalized turbidity above 0.3 are colored in red.

**Figure S6.**
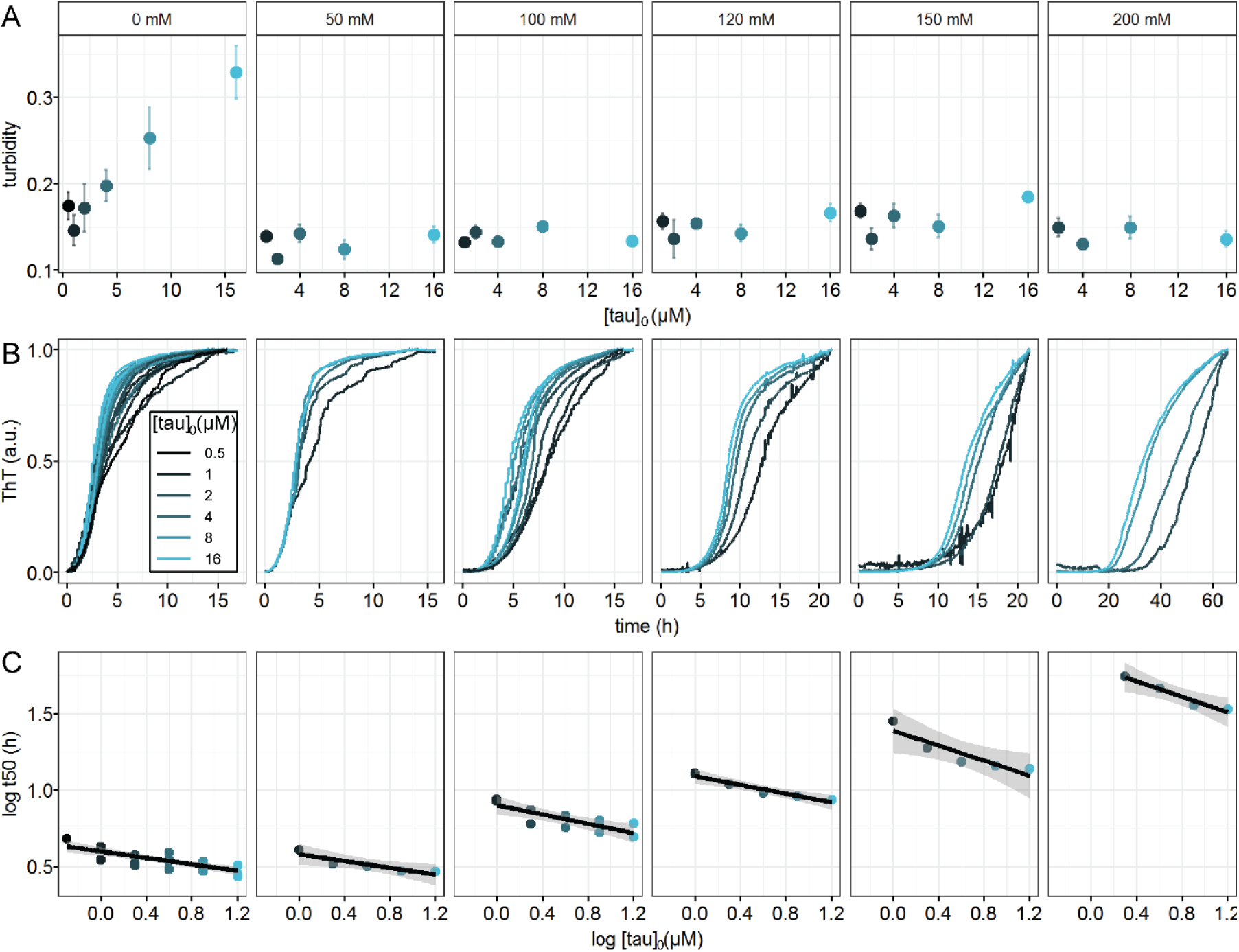
Effects of LLPS on tau-heparin aggregation monomer dependence. **A.** Turbidity of tauSS-heparin mixture at varying initial tau concentration, [tau]_0_. Ratio of heparin was fixed at 1.7 *μ*g/mL heparin per 1 *μ*M tau. **B**. Normalized ThT fluorescence of tauSS-heparin mixture at varying [tau]_0_ and [NaCl]. Data were fit with sigmoid function to extract t_1/2_ (see Materials and Methods). **C.** log-log plot of t_1/2_ vs [tau]_0_. Solid lines are linear regression of the data points. Shade areas show confidence interval of the regression. Slopes of the fit, or *γ*, are −0.11 ± 0.02, −0.11 ± 0.03, −0.15 ± 0.04, −0.14 ± 0.02, −0.24 ± 0.06, −0.26 ± 0.04, for each panel from left to right respectively. R^2^ of the regression are 0.64, 0.82, 0.68, 0.94, 0.84, 0.95, for each panel from left to right respectively.

**Figure S7.**
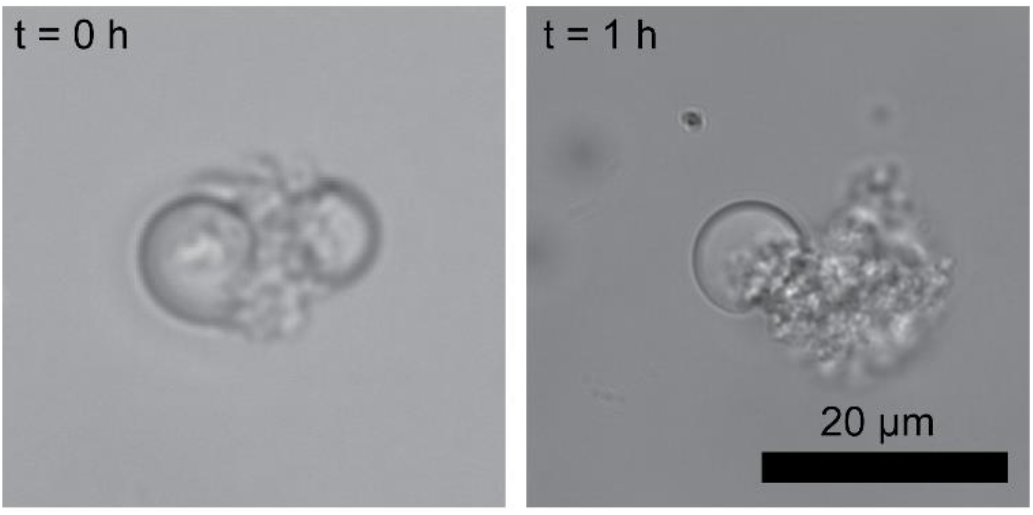
Direct contacts of tau-RNA droplets with tau aggregates. 20 μM tauSSP301L was mixed with 60 μg/mL poly(U) RNA in the presence of 1 μM seed (prepared as below). Samples were loaded onto a microscope slide with glass cover slide and incubated at room temperature for 1 hour before images were taken. Seed was prepared by incubating 20 μM tauSSP301L with 5 μM heparin overnight at 37 °C.

